# Nanoscale manipulation of membrane curvature for probing endocytosis in live cells

**DOI:** 10.1101/122275

**Authors:** Wenting Zhao, Lindsey Hanson, Hsin-Ya Lou, Matthew Akamatsu, Praveen D. Chowdary, Francesca Santoro, Jessica R. Marks, Alexandre Grassart, David G. Drubin, Yi Cui, Bianxiao Cui

## Abstract

Clathrin-mediated endocytosis (CME) involves nanoscale bending and inward budding of the plasma membrane, by which cells regulate both the distribution of membrane proteins and the entry of extracellular species^1,2^. Extensive studies have shown that CME proteins actively modulate the plasma membrane curvature^1,3,4^. However, the reciprocal regulation of how plasma membrane curvature affects the activities of endocytic proteins is much less explored, despite studies suggesting that membrane curvature itself can trigger biochemical reactions^5-8^. This gap in our understanding is largely due to technical challenges in precisely controlling the membrane curvature in live cells. In this work, we use patterned nanostructures to generate well-defined membrane curvatures ranging from +50 nm to -500 nm radius of curvature. We find that the positively curved membranes are CME hotspots, and that key CME proteins, clathrin and dynamin, show a strong preference toward positive membrane curvatures with a radius < 200 nm. Of ten CME related proteins we examined, all show preferences to positively curved membrane. By contrast, other membrane-associated proteins and non-CME endocytic protein, caveolin1, show no such curvature preference. Therefore, nanostructured substrates constitute a novel tool for investigating curvature-dependent processes in live cells.

---

Membrane curvature is no longer seen merely as a passive feature of membranes, but has emerged as a highly active player in regulating protein activities^9,10^. To date, how membrane curvature affects protein binding and activity has been primarily studied using *in vitro* systems, such as supported lipid bilayers and lipid vesicles^8,11,12^. However, these findings need to be validated in live cells, which in the context of endocytosis requires controlling plasma membrane curvature at the scale of tens to hundreds of nanometer radius. This is a challenging task as the plasma membrane is a dynamic and complex system containing hundreds of different lipid and protein components. A recent study has shown that the plasma membrane can be deformed on nanoscale conical structures (nanocones), which triggers the recruitment of curvature-sensing N-BAR proteins^13^. However, as nanocones were variables in sizes and densely packed^14^, the extent of nanocone-induced membrane curvature was not well controlled and not individually discernible under an optical microscope^13^. Conversely, we and others have shown that vertically-aligned nanopillars that are evenly spaced and uniform in size can induce conformal plasma membrane wrapping in mammalian cells^15-17^. Based on these studies, we hypothesize that the shape and geometry of vertically-aligned nanostructures can be used to control local membrane curvature in live cells (**Fig. 1a**). In this work, we demonstrate that vertically-aligned nanostructure arrays generate well-defined membrane curvature, which we exploit to probe how membrane curvature affects CME activities.

**Figure 1.**
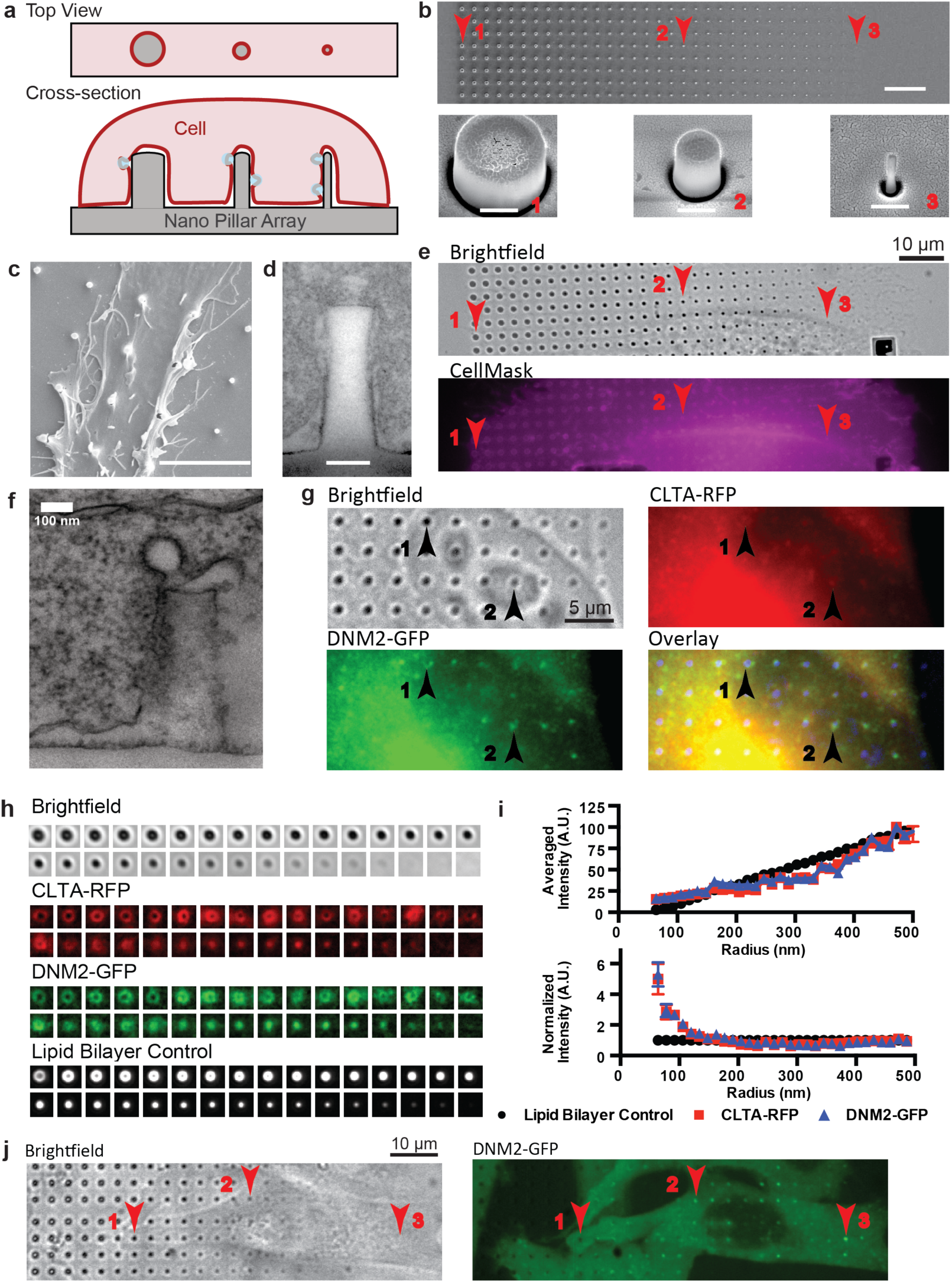
Vertical nanopillars generate well-defined membrane curvatures that induce local accumulation of endocytic proteins. **a.** Schematic illustration of nanopillars with different radii (grey block) deforming the cell membrane to generate different membrane curvatures (red line). **b.** An SEM image of a gradient nanopillar array with 700 nm height, 3 μm center-to-center distance between nanopillars, and radii ranging from 500 nm (left) to 50 nm (right) with 14 nm increment. Scale bar: 10 μm. Bottom row: zoomed-in SEM images of individual nanopillars as indicated by corresponding locations in the top row (red with arrow). Scale bar: 400 nm. **c.** An SEM image showing a SK-MEL-2 cell deforming on nanopillars. Scale bar: 5 μm. **d.** A TEM image showing the cell membrane wrapping around a nanopillar. Scale bar: 100 nm. **e.** The plasma membrane stained with CellMask™ DeepRed shows increased membrane signal at nanopillar locations. Red arrows indicate locations of nanopillars with different radius. Scale bar: 10 μm. **f.** A TEM image of the plasma membrane wrapping around a nanopillar captures a clathrin-coated pit. Scale bar: 100 nm. **g.** Immunostaining of clathrin and dynamin2 in h*CLTA*^EN^/h*DNM2*^EN^ cells shows accumulation of clathrin and dynamin2 at nanopillar locations. In the overlay image, the inverted brightfield image is converted to blue (bottom right). Scale bar: 10 μm. Numbered black arrows indicate locations of nanopillars with different radii. **h.** Quantification of the clathrin, dynamin2 and membrane signals at nanopillars of 32 different radii. Signals from many nanopillars of the same radius are averaged to obtain the averaged image. Averaged images were sorted from largest (top left) to smallest radius (bottom right). Four imaging channels are shown from top to bottom: bright field, clathrin, dynamin2 and lipid bilayer control for surface area increment. **i.** Intensities of clathrin-RFP, dynamin2-GFP and lipid membrane all increase with nanopillar radius (top graph). After normalizing against the membrane intensities, it is clear that the clathrin/membrane and dynamin2/membrane ratios increase significantly when the nanopillar radius is less than 200 nm (bottom graph). Each data point is an average over 90-191 nanopillars with error bar representing standard error of the mean. Detailed statistic analysis is in **Suppl. Table S5. j.** The time-averaged image of a 4 min movie of dynamin2-GFP demonstrates that dynamin2-GFP exhibits strong preference to sharp nanopillars, but much less to large radius nanopillars in the same cell (Arrow 1 *vs*. Arrow 3). Scale bar: 10 μm.

To demonstrate that membrane curvature can be controlled using vertically-aligned nanopillars, we engineered a gradient array of SiO_2_ nanopillars with different radii from 50 nm to 500 nm with a 14 nm increment (scanning electron microscopy (SEM) see **Fig. 1b**).The size variations are within 10% of the nominal value (**Suppl. Fig. S1, Table S1-4**). The area along the side of each nanopillar is about 10 times the area of the top, so we primarily considered the membrane curvature along the side, a positive curvature denoted by the nanopillar radius. These curvatures cover the mid range of endocytic curvature progression (from flat to ˜50 nm radius). SK-MEL-2 cells cultured on a poly-L-lysine coated substrate bend their membrane on nanopillars, as seen in the SEM image in **Fig. 1c**. This was substantiated by the cross-section visualization using TEM (**Fig. 1d**) and focused ion beam and scanning electron microscopy (FIB-SEM) (**Suppl. Fig. S2**). The plasma membrane wraps around nanopillars with an average gap distance of 21.6± 14.1 nm (mean ± s.d.), consistent with previous findings^15^. Membrane wrapping around nanopillars of all sizes was further confirmed by membrane staining (CellMask™ Deep Red, **Fig. 1e**), where rings appeared on large nanopillars and dots appeared on small nanopillars below the diffraction limit. The fluidity of the membrane around nanopillars was verified to be similar to that of flat areas using fluorescence recovery after photobleaching (**Suppl. Fig. S3, Movie S1**). These results confirm that vertically-aligned nanopillars can induce well-defined plasma membrane curvatures.

In TEM (**Fig. 1f**) and FIB-SEM (**Suppl. Fig. 2e**), we captured a few endocytic events as shown by the characteristic clathrin-coated pit profiles, indicating that CME does occur on nanopillar-curved membranes. To investigate CME further, we used gradient nanopillar arrays to examine how membrane curvature affects the distribution of two key CME proteins, clathrin and dynamin^18,19^. Immunostaining clearly showed increased clathrin and dynamin signals at nanopillars (arrows, **Fig. 1g**). To account for cell-to-cell variations, we averaged signals from over 2000 nanopillars in 41 cells to obtain averaged fluorescence images of clathrin and dynamin at each nanopillar radius (**Fig. 1h**). As a control for the surface area, we also measured the fluorescence signal of a supported lipid membrane (**Fig. 1h**, bottom panel), a membrane associated protein (GFP-CAAX), and CellMask staining on the same array (**Suppl. Fig. S4**). The fluorescent signals for the membrane, clathrin and dynamin all increased with increasing nanopillar radius (**Fig. 1i**, top). By normalizing the intensities of clathrin and dynamin against that of the membrane (**Fig. 1i**, bottom), we found that the ratios of clathrin/membrane and dynamin/membrane are relatively constant for radii > 200 nm. However, for radii < 200 nm, both ratios increase significantly, indicating a clear preference of clathrin and dynamin for curvature radius < 200 nm. This strong preference for high curvature is also evident in single cells. In an averaged image from a 4-min movie (**Fig. 1j** shows), dynamin2-GFP exhibits stronger signals at small-radius nanopillars than at large-radius nanopillars in the same cell (Arrow 3 *vs*. Arrow 1 in **Fig. 1j**), despite their smaller membrane area.

The preferential accumulation of proteins on highly curved membranes is more compellingly demonstrated by nanostructures that generate a local combination of different curvatures. As illustrated in **Fig. 2a**, a single nanobar structure locally induces two different membrane curvatures – high curvature at the two ends and zero curvature in the middle. **Fig. 2b** shows an SEM image of a nanobar array. When cells were cultured on the nanobar array, the plasma membrane was found to wrap evenly around the bars, outlining the nanobar shape as shown by CellMask staining (**Fig. 2c**). To probe protein dynamics, genome-edited SK-MEL-2 cells (hCLZA^EN^/hDNM2^EN^)^19^ that express clathrin-RFP and dynamin2-GFP were cultured on nanobar arrays. Unlike the CellMask staining, both clathrin and dynamin2 showed strong preferences for the highly curved ends of the nanobars and very little accumulation with the middle region (**Fig. 2d**). The averaged images from over 167 nanobars (**Fig. 2e**) more clearly illustrated the strong accumulation of clathrin and dynamin at nanobar ends (**Fig. 2f**). Kymograph analysis of time-lapse movies demonstrated that both clathrin and dynamin2 were dynamic and had strong preference for nanobar ends (**Fig. 2g**, **Suppl. Fig. S5, Movie S2)**. Endocytic events, as evidenced by the characteristic appearance of dynamin2 near the end of the clathrin lifetime^19^, are abundant at nanobar ends, while very few events are observed in the middle of the same nanobars. Similarly, the endocytic adaptor protein AP2 (AP2-RFP) also shows preferred accumulation on nanopillars with the appearance of dynamin2-GFP at the end of the AP2 lifetime (**Suppl. Fig. S6, Movie S3**).

**Figure 2.**
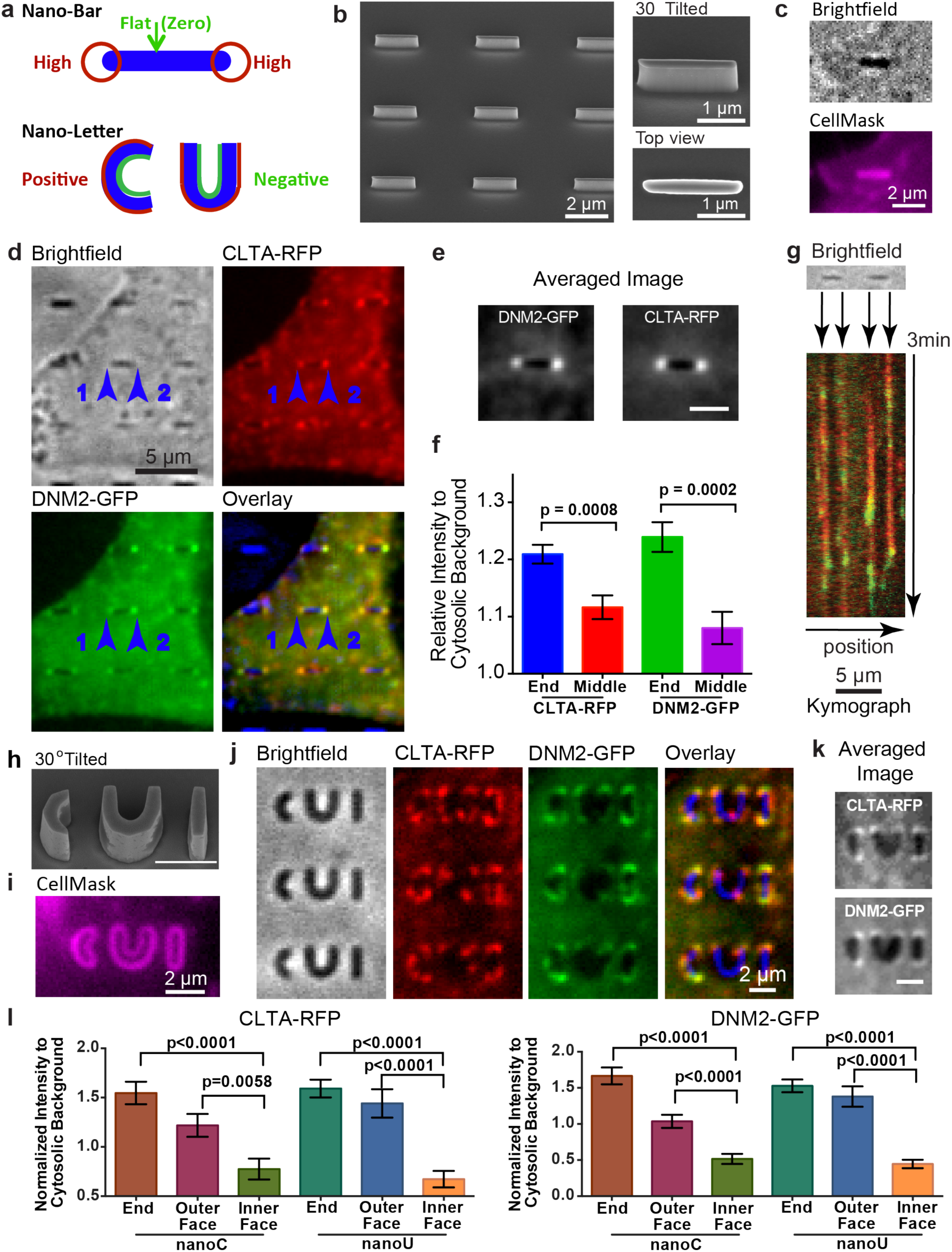
Engineered 3D nanostructures for versatile control of membrane curvatures and endocytic protein accumulations. **a.** Schematic illustration of nanobar and nano-letter design for inducing a range from flat membranes to high membrane curvatures (top, nanobar) and combined positive and negative membrane curvatures (bottom, nanoC and nanoU) by the same nanostructure. **b.** An SEM image of a nanobar array showing individual nanobars of 150 nm width, 2 μm length, 1 μm height and 5 μm pitch(left, scale bar: 2 μm.). **c.** When SK-MEL-2 cells were cultured on nanobar arrays, CellMask™ DeepRed staining demonstrated that the plasma membrane wrapped evenly around the nanobar. Scale bar: 2 μm. **d.** Averaged fluorescence images of a 4 min movie shows that clathrin and dynamin2 prefer the two highly curved ends of nanobars as compared with the sidewalls. This is in sharp contrast to the membrane staining shown in c. In the overlay image, bright field images of nanobars are shown in blue. Scale bar: 5 μm. **e.** Averaged image of clathrin and dynamin2 distribution on 167 nanobars. Scale bar: 2 μm. **f.** The quantitative values of fluorescence intensity at nanobar ends and nanobar center with the cytosolic background as reference. Error bar represents standard error of the mean. Statistic analysis shows significant difference between nanobar end and nanobar center (p value of unpaired t-test: 0.0008 for Clathrin-RFP and 0.0002 for Dynamin2-GFP). **g.** Kymograph plots of clathrin-RFP (red) and dynamin2-GFP (green) on two adjacent nanobars show the appearance of a dynamin2 peak near the end of the clathrin segment, which is characteristic of clathrin-mediated endocytosis. The dynamic events are observed at the ends of nanobars, with very little signal along the side walls or between nanobars. More kymographs on a larger area with more adjacent nanobars are in **Suppl. Fig. S5. h.** An SEM image of the nanoCUI structure in 30° tilted view. Scale bar: 2 μm. **i.** CellMask™ DeepRed staining of cells on nanoCUI array shows that the cell membrane wrapped around both the inside and the outside surfaces of the nanoC and nanoU structures. Scale bar: 2 μm. **j.** Averaged fluorescence images of a 4 min movie show that clathrin and dynamin2 prefer positive membrane curvatures at the ends and the outer surfaces of nanoC and nanoU, with much less protein signal on the inner surfaces. **k.** Averaged images of clathrin-RFP (top) and Dynamin2-GFP (bottom) on 51 pairs of nanoC and nanoU clearly show preferred accumulation on the ends and their outer faces of both nanoC and nanoU comparing to the inner faces. Scale bar: 2 μm. **l.** The quantified fluorescence intensity (in k) on the end, the outer face, and the inner face of both nanoC (left) and nanoU (right) with the cytosolic background as reference. Error bar represents standard error of the mean. Statistic analysis shows significant difference at the end/the outer face *vs*. the inner face of both nanoC and nanoU. The p value of unpaired t-test for each pair is indicated on the plot.

In addition to probing membrane curvatures of different magnitudes, we also probed curvatures of opposite signs. We engineered nanoC and nanoU to induce positive curvature at the outer face (**Fig. 2a**, red line) and negative curvature at the inner face (**Fig. 2a**, green line). In order to distinguish the outer and inner faces by optical microscopy, we made nanoC and nanoU structures of 500 nm width (**Fig. 2h**). Membrane staining by CellMask clearly illustrates the inner and the outer faces and confirms that nanoC and nanoU induced both positive and negative membrane curvatures (**Fig. 2i**). When hCLTA^EN^IhDNM2^EN^ cells were cultured on nanoC and nanoU, both clathrin and dynamin2 exhibited accumulation on the outer surface of the nanostructures with positive membrane curvature. By contrast, very little protein signal appeared on the negative curvature side (**Fig. 2j**). The averaged images from over 50 nanoCUI structures show that both clathrin and dynamin prefer high curvature locations at the ends of nanoC, nanoU and nanoI (**Fig. 2k**). For nanoC and nanoU, the fluorescence intensities at the outer surface are statistically higher than on the inner surface (**Fig. 2l**). These results demonstrate that endocytosis, a process of positive curvature progression, is favored by preformed positive, but not negative, curvature.

Besides clathrin and dynamin2, many other proteins and lipids participate in CME^20^ (schematic see **Fig. 3a**). We examined the curvature preference of twelve proteins on nanobar arrays, including those with curvature-sensing domains (e.g., epsin with an ENTH domain^21^ and amphiphysin with an N-BAR domain^22^) and those not reported to have curvature-sensing domains (e.g., AP2^23^). The high magnification fluorescence images in **Fig. 3 b-o** (green color images) show the spatial distributions of different proteins on six nanobars (the corresponding full images are shown in **Suppl. Fig. S7-S8**). For each protein, the averaged image over hundreds of nanobars is shown in grey-scale and the protein intensity is plotted along the length of the nanobar (**Fig. 3 b-o**). Their curvature preference was evaluated by the ratio of the intensity at the nanobar-end *vs*. nanobar-center (**Fig. 3p, Suppl. Table S6**).

**Figure 3.**
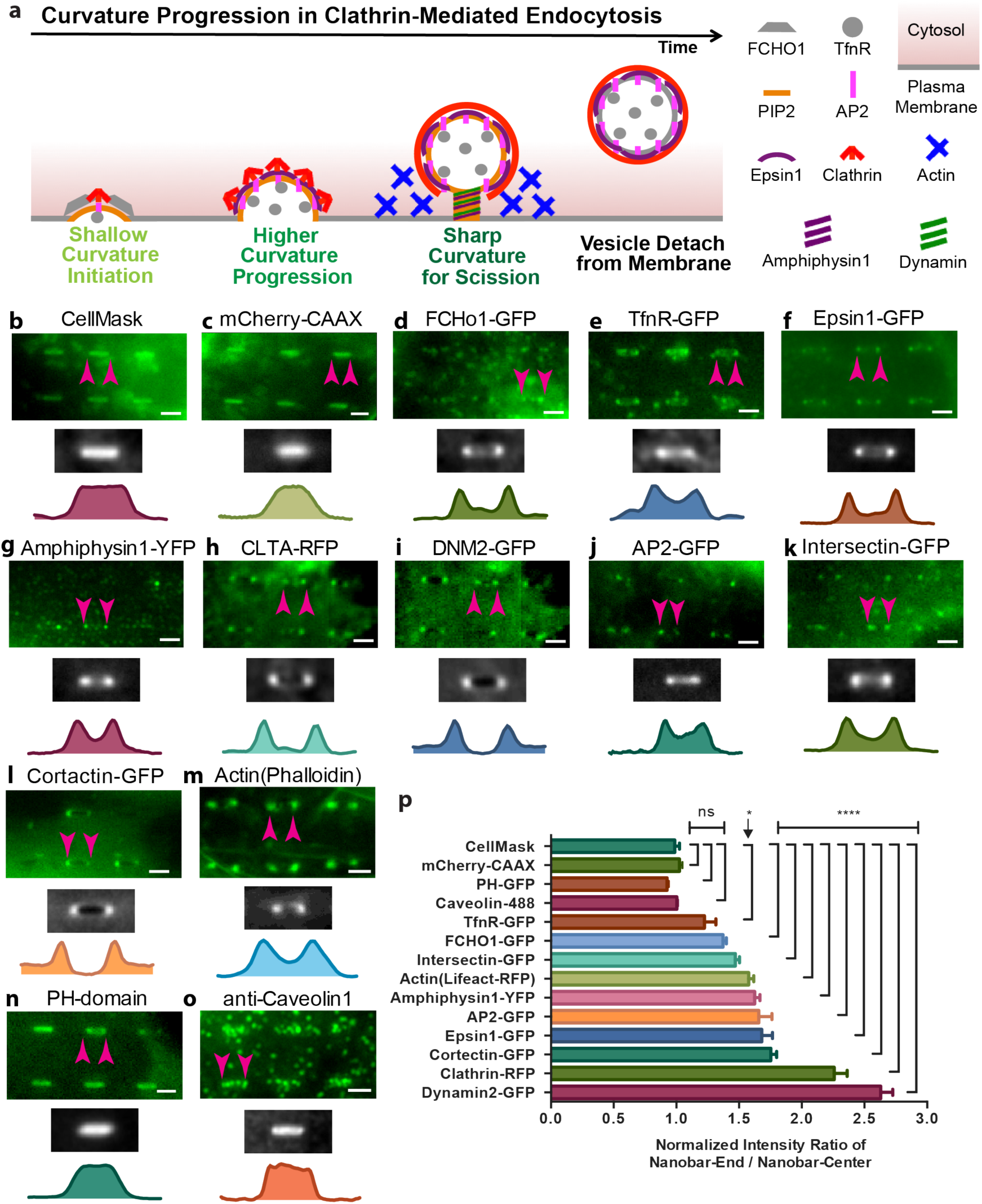
Probing curvature sensitivity of various endocytic proteins using nanobar arrays. **a.** Schematic illustration of proteins involved in different stages of clathrin-mediated endocytosis. The high magnification fluorescence images in **b-o** (green color images, scale bar: 2 μm) show the distributions of different proteins on six nanobars. The corresponding large images can be found in the supplementary information (**Fig. S6 and S7**). The averaged image of hundreds of nanobars and the intensity along the length of nanobars are shown beneath the fluorescent images. The spatial distributions of CellMask™ DeepRed staining the plasma membrane (**b**), and membrane-associated protein mCherry-CAAX (**c**) are distributed relatively evenly along the entire length of the nanobars. Eight other endocytic components involved in the clathrin-dependent endocytosis including epsin1-GFP (**f**), amphiphysin1-YFP (**g**), clathrin-RFP (**h**), dynamin2-GFP (**i**), AP2-GFP (**j**), intersectin (**k**), cortactin (**l**) and actin (**m**) show strong bias toward the ends of the nanobars with little accumulation along the sidewalls. In comparison, FCHol-GFP (**e**), and transferrin receptor (TfnR-GFP) (**f**) show less preference to the nanobar ends than those above. On the other hand, PIP2 probed by PH-GFP (**n**) and caveolin1 (**o**), an essential component of the caveolin-dependent endocytosis, is evenly distributed along the entire length of nanobars. All the nanobars are 2 μm in length. p. Distribution of 14 different proteins/dyes on nanobars quantified by normalized intensity ratio of nanobar-end to nanobar-center. For each protein, the distribution is measured by averaging over 56-2072 nanobars. Error bar represents standard error of the mean. Statistical significance of each protein *vs*. CellMask was evaluated by unpaired t-test with Welch’s correction (details see **Suppl. Table S6**). p-value: **** < 0.0001, * < 0.05, ns >0.05.

As a reference for protein distribution, the plasma membrane was probed by CellMask (**Fig. 3b**) and a plasma membrane-associated protein mCherry-CAAX (**Fig. 3c**), both were homogeneous along the length of the nanobar. In sharp contrast, all 10 CME proteins showed spatial preference for the highly curved ends of nanobars (**Fig. 3. d-m**). Proteins involved in the early stages of CME such as FCHo-1-GFP^24^ (**Fig. 3d**) and transferrin receptor (TfnR-GFP)^25^ (**Fig. 3e**) showed moderate preference to nanobar ends. Proteins involved in the later stages of CME showed much stronger preference, including epsin1-GFP (**Fig. 3f,**), amphiphysin1-YFP (**Fig. 3g**,), CLTA-RFP (**Fig. 3h**,), DNM2-GFP (**Fig. 3i**,), AP2-GFP (**Fig. 3j**,), intersectin (**Fig. 3k**,), cortactin (**Fig. 3l**,), and actin (**Fig. 3m**). Their averaged images show a characteristic dumbbell shape and the intensity plots indicate two pronounced peaks at the two ends of the nanobars (**Fig. 3f-m**). Of all the CME proteins, AP2, intersectin, cortactin and actin are not reported to have curvature sensitive-domains. Their accumulation at nanobar ends strongly indicates that both curvature sensitive and non-sensitive CME proteins co-assemble at highly curved membrane for CME progression.

Although CME proteins show strong preference to highly curved membrane, a signaling lipid involved in the CME process, phospholipid phosphatidylinositol 4,5-biphosphate (PIP2)^26,27^, did not show such preference. PIP2 was probed using the PH domain of PLC-delta (PH-GFP) (**Fig. 3n)**, which showed uniform distribution along the nanobar similar to CellMask. We also examined the curvature sensing of caveolin1, an essential protein in caveolin-dependent endocytosis^28^. As shown in **Fig. 3o**, caveolin1 formed puncta evenly distributed along the entire length of nanobars with no preference toward the nanobar ends. This indicates that caveolin-dependent and clathrin-mediated endocytosis are affected differently by membrane curvature.

In addition to probing the spatial preference of CME proteins for membrane curvatures, we also investigated their dynamics. We compared the occurrence frequency of dynamin2 spots on nanopillars of 150 nm radius *vs*. on flat areas. Line kymographs crossing several nanopillars showed clear segments of dynamin2 traces aligned with nanopillars (**Fig. 4a**, blue lines), while very few dynamin2 events on adjacent flat areas (**Fig. 4a**, red lines). The rapid appearance and disappearance of a dynamin2 spot is a signature of an endocytic vesicle scission event^19^. A spatial map of dynamin2 events clearly shows hotspots for endocytosis at nanopillars (**Fig. 4b**). After normalizing for the membrane area, there is significantly more occurrence of dynamin2 clusters on nanopillars than on flat membranes (**Fig. 4b**), indicating that pre-curved membranes are preferred CME sites.

**Figure 4.**
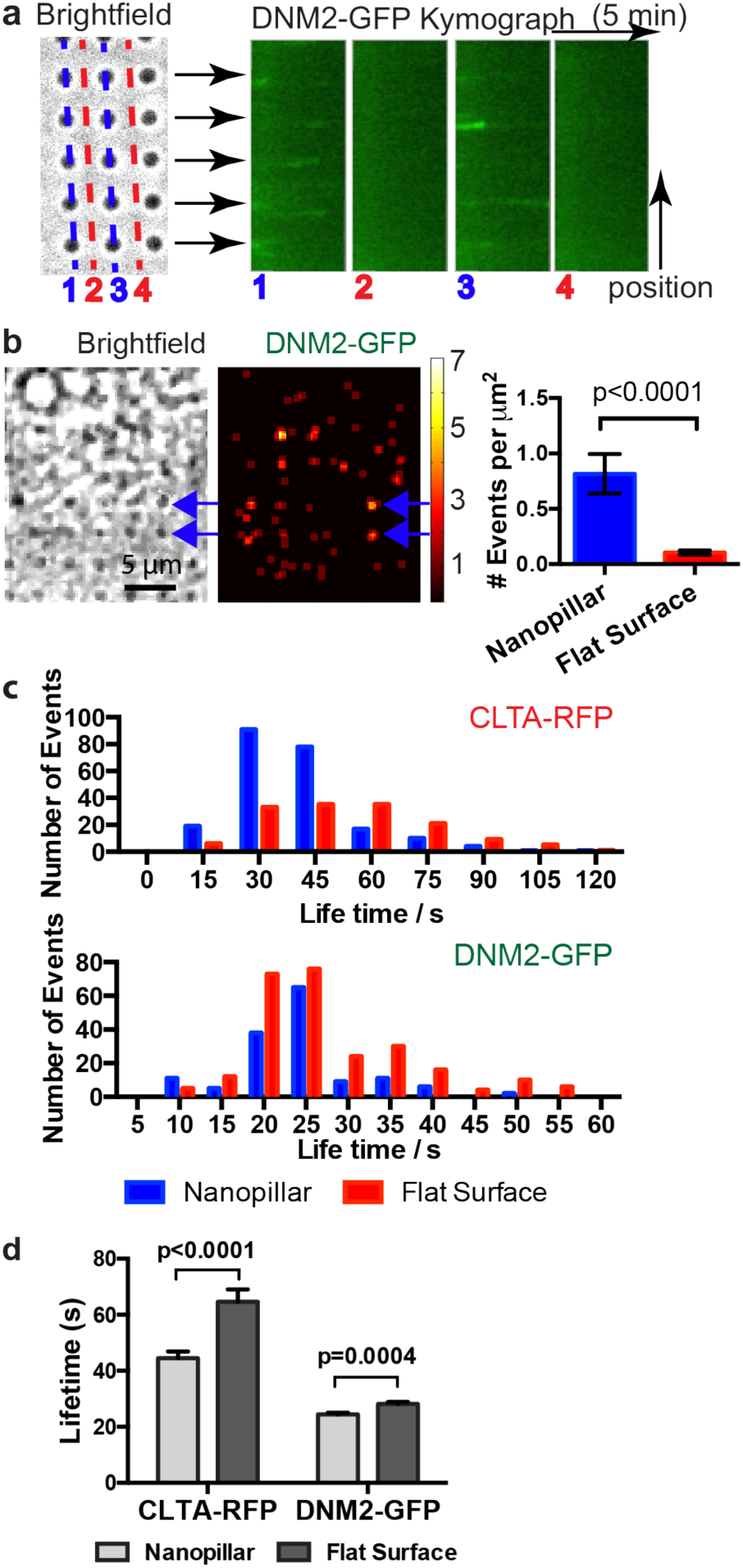
Pre-curved membranes are preferred sites for endocytosis. **a.** Kymograph plots of dynamin2-GFP along lines of nanopillars show repeated appearance and disappearance of dynamin2 spots at nanopillar locations (line 1 and 3). Similar kymograph plots along lines between nanopillars show very little signal of dynamin2-GFP (line 2 and 4). **b.** Spatial mapping of the occurrence frequency of dynamin2-GFP puncta shows hot spots for endocytosis at nanopillar locations. We measured 65 dynamin2 blinking events on 75 nanopillars in 6 minutes. Nanopillar area is calculated to be 1 μm^2^ each. From the same set of movies, we measured 62 dynamin2 blinking events on 75 flat areas, 8 μm^2^ each. After normalizing against the calculated membrane areas, the endocytic event occurrence on curved membrane at nanopillars is significantly higher than that on flat. Error bars represent standard error of the mean and the P value of Kolmogorov–Smirnov test is < 0.0001. **c.** Lifetime distribution of clathrin-RFP (up) and dynamin2-GFP (bottom) puncta on nanopillars (blue) and on flat areas (red). For clathrin, 226 events were measured on nanopillars and 154 events were measured on flat areas. For dynamin2, 147 events were measured on nanopillars and 260 events were measured on flat areas. For comparison between nanopillar and flat, Kolmogorov–Smirnov test gives p-value of 0.0082 for dynamin2-GFP and <0.0001 for clathrin-RFP. The differences are significant. **d.** The average lifetimes of clathrin-RFP and dynamin2-GFP appear to be decreased on nanopillars (light grey) compared with on flat areas (dark grey). Unpaired t-test with equal SD gives p-value of 0.0004 for dynamin2-GFP and <0.0001 for clathrin-RFP. The differences are significant.

Besides frequency, we also measured the lifetime of dynamin2 and clathrin spots on nanopillars *vs*. on flat areas (**Suppl. Movie S4).** The dynamin2 lifetime distribution (**Fig. 4c**) shows two apparent populations on flat areas, one around 20 s and the other around 35 s. On nanopillars, the dynamin2 lifetime distribution also shows similar peaks, but the population with longer lifetimes is slightly decreased. The average lifetime of dynamin2 is slightly shorter on nanopillars (24±0.6 s *vs*. 28±0.7 s, mean±SEM) (**Fig. 4d**). On the other hand, the lifetime for clathrin spots appears to decrease significantly on nanopillars as compared to flat areas (45±2.3 s vs. 65±4.3 s, mean±SEM). It is plausible that pre-curved membrane reduces the energy barrier for the membrane bending and thus facilitates faster clathrin coat assembly. However, we note that the lifetime measurement may miss some long clathrin events as multiple endocytosis events sometimes overlap on nanopillars.

Our results demonstrate that CME proteins show spatial preference to highly positively curved over flat or negatively curved membranes in live cells. We also confirm that membrane curvature can directly modulate the biochemical activities of CME, as previously proposed^5^. Interestingly, actin also shows a strong preference for highly curved membranes. As actin and its regulators are involved in diverse cellular functions^29,30^, these results suggest that membrane curvature may affect many other cellular processes in addition to endocytosis. The ability to precisely manipulate membrane curvature in live cells opens up new avenues to study the influence of membrane curvature on a variety of cellular processes.

## Methods

Methods and associated references are described in the supplementary information.

## Author Contribution

W.Z., B.C., Y.C., and D.G.D conceived the study and designed the experiment. W.Z. fabricated the nanostructure substrates, and performed most of experiments. L.H. performed TEM measurements. F.S. conducted the FIB/SEM characterization. H.Y.L. performed most of the endocytic protein test on nanobar arrays and the quantification and statistic analysis. W.Z., P.C. and B.C. developed the Matlab code for the dynamic analysis. W.Z. analyzed the most of the data. M.A. analyzed the AP2/Dynamin2 movies. A.G., and J.M provided and characterized the genome-edited cell line. W.Z. and B.C. wrote the manuscript. All the authors discussed the results and commented on the manuscript.

## Acknowledgement

We thank Dr. Yansong Miao and Dr. Sun Hae Hong of the Drubin group in U.C. Berkeley for valuable discussion as well as helpful comments on genome-edited cell lines and endocytic lifetime analysis, Prof. Kang Shen in Stanford for generous support on spinning disk confocal microscopy, Dr. Milos Galic of the Tobias Meyer group in Stanford for suggestions and Amphiphysin1-YFP plasmid, Dr. Shunling Guo of the Bianxiao Cui group for constructing mCherry-CAAX plasmid, as well as Allister McGuire, Dr. Chong Xie and Dr. Ziliang Lin of the Bianxiao Cui group in Stanford for advice and help on the nanostructure fabrication. We also thank Qunxiang Ong and Luke Kaplan of Bianxiao Cui group for comments on the manuscript. Fabrication and characterization of nanostructures were conducted in Stanford Nanofabrication Center and Stanford Nano Shared Facilities (SNSF). Spinning disk confocal with perfect focus for lifetime measurement was conducted in Cell Science Imaging Facility (CSIF) at Stanford University. This work was supported by the NSF (CAREER award no. 1055112), the NIH (grant no. NS057906), a Searle Scholar award, a Packard Science and Engineering Fellowship (to B.C.), NIH fellowship 1F32 GM113379 (to J.R.M.) and the NIH (R01 GM65462) (to D.G.D.).

## References

1. McMahon, H. T. & Boucrot, E. Molecular mechanism and physiological functions of clathrin-mediated endocytosis. Nat Rev Mol Cell Biol 12, 517–533 (2011).

2. Di Fiore, P. P. & Zastrow, von, M. Endocytosis, signaling, and beyond. Cold Spring Harb. Perspect. Biol. 6, (2014).

3. Johannes, L., Wunder, C. & Bassereau, P. Bending ‘on the rocks’‐‐a cocktail of biophysical modules to build endocytic pathways. Cold Spring Harb. Perspect. Biol. 6, a016741–a016741 (2014).

4. Kirchhausen, T., Owen, D. & Harrison, S. C. Molecular Structure, Function, and Dynamics of Clathrin-Mediated Membrane Traffic. Cold Spring Harb. Perspect. Biol. 6, a016725–a016725 (2014).

5. McMahon, H. T. & Gallop, J. L. Membrane curvature and mechanisms of dynamic cell membrane remodelling. Nature 438, 590–596 (2005).

6. Liu, J., Sun, Y., Drubin, D. G. & Oster, G. F. The mechanochemistry of endocytosis. PLoS Biol 7, e1000204 (2009).

7. Galic, M. et al. Dynamic recruitment of the curvature-sensitive protein ArhGAP44 to nanoscale membrane deformations limits exploratory filopodia initiation in neurons. eLife 3, e03116 (2014).

8. Larsen, J. B. et al. Membrane curvature enables N-Ras lipid anchor sorting to liquid-ordered membrane phases. Nat Chem Biol 11, 192–194 (2015).

9. Epand, R. M., D’Souza, K., Berno, B. & Schlame, M. Membrane curvature modulation of protein activity determined by NMR. Biochim. Biophys. Acta 1848, 220–228 (2015).

10. Iversen, L., Mathiasen, S., Larsen, J. B. & Stamou, D. Membrane curvature bends the laws of physics and chemistry. Nat. Cell Bio. 11, 822–825 (2015).

11. Wu, M. et al. Coupling between clathrin-dependent endocytic budding and F-BAR-dependent tubulation in a cell-free system. Nat. Cell Bio. 12, 902–908 (2010).

12. Lee, I.-H., Kai, H., Carlson, L.-A., Groves, J. T. & Hurley, J. H. Negative membrane curvature catalyzes nucleation of endosomal sorting complex required for transport (ESCRT)-III assembly. Proc. Natl. Acad. Sci. U.S.A. 112, 15892–15897 (2015).

13. Galic, M. et al. External push and internal pull forces recruit curvature-sensing N-BAR domain proteins to the plasma membrane. Nat. Cell Bio. 14, 874–881 (2012).

14. Jeong, S., McDowell, M. T. & Cui, Y. Low-Temperature Self-Catalytic Growth of Tin Oxide Nanocones over Large Areas. ACS Nano 5, 5800–5807 (2011).

15. Hanson, L., Lin, Z. C., Xie, C., Cui, Y. & Cui, B. Characterization of the cell-nanopillar interface by transmission electron microscopy. Nano Lett. 12, 5815–5820 (2012).

16. Mumm, F., Beckwith, K. M., Bonde, S., Martinez, K. L. & Sikorski, P. A transparent nanowire-based cell impalement device suitable for detailed cell-nanowire interaction studies. Small 9, 263–272 (2013).

17. Santoro, F. et al. Interfacing electrogenic cells with 3D nanoelectrodes: position, shape, and size matter. ACS Nano 8, 6713–6723 (2014).

18. Avinoam, O., Schorb, M., Beese, C. J., Briggs, J. A. G. & Kaksonen, M. Endocytic sites mature by continuous bending and remodeling of the clathrin coat. Science 348, 1369–1372 (2015).

19. Doyon, J. B. et al. Rapid and efficient clathrin-mediated endocytosis revealed in genome-edited mammalian cells. Nat. Cell Bio. 13, 331–337 (2011).

20. Taylor, M. J., Perrais, D. & Merrifield, C. J. A High Precision Survey of the Molecular Dynamics of Mammalian Clathrin-Mediated Endocytosis. PLoS Biol 9, e1000604 (2011).

21. Ford, M. G. J. et al. Curvature of clathrin-coated pits driven by epsin. Nature 419, 361–366 (2002).

22. Peter, B. J. BAR Domains as Sensors of Membrane Curvature: The Amphiphysin BAR Structure. Science 303, 495–499 (2004).

23. AP2 controls clathrin polymerization with a membrane-activated switch. 345, 459–463 (2014).

24. Henne, W. M. et al. FCHo Proteins Are Nucleators of Clathrin-Mediated Endocytosis. Science 328, 1281–1284 (2010).

25. Dautry-Varsat, A., Ciechanover, A. & Lodish, H. F. pH and the recycling of transferrin during receptor-mediated endocytosis. Proc. Natl. Acad. Sci. U.S.A. 80, 2258–2262 (1983).

26. Di Paolo, G. & De Camilli, P. Phosphoinositides in cell regulation and membrane dynamics. Nature 443, 651–657 (2006).

27. Posor, Y. et al. Spatiotemporal control of endocytosis by phosphatidylinositol-3,4-bisphosphate. Nature 499, 233–237 (2013).

28. Chaudhary, N. et al. Endocytic Crosstalk: Cavins, Caveolins, and Caveolae Regulate Clathrin-Independent Endocytosis. PLoS Biol 12, e1001832 (2014).

29. Grassart, A. et al. Actin and dynamin2 dynamics and interplay during clathrin-mediated endocytosis. J. Cell Biol. 205, 721–735 (2014).

30. Dominguez, R. & Holmes, K. C. Actin Structure and Function. Annu. Rev. Biophys. 40, 169–186 (2011).

